# Host genotype and sex shape influenza evolution and defective viral genomes

**DOI:** 10.1101/2025.02.26.638946

**Authors:** Rodrigo M. Costa, Lehi Acosta-Alvarez, Kaili Curtis, Kort Zarbock, Justin Kelleher, Bhawika S. Lamichhane, Andrew L. Valesano, William J. Fitzsimmons, Adam S. Lauring, Jon Seger, Frederick R. Adler, Wayne K. Potts

**Affiliations:** School of Biological Sciences, University of Utah, Salt Lake City, UT, USA; Department of Neurology and Neurological Sciences, Stanford University School of Medicine, Stanford, CA, USA; Department of Internal Medicine and Department of Microbiology and Immunology, University of Michigan, Ann Arbor, MI, USA

## Abstract

Viral evolution during initial pandemic waves favors mutations that enhance replication and transmission over antigenic escape. Host genotype and sex strongly shape this early adaptation, yet their individual and combined effects remain unclear. We experimentally adapted influenza A virus to male and female BALB/c and C57BL/6 mice, generating 28 independent lineages, and employed a novel “rolling sphere” approach to identify mutational hotspots in three-dimensional protein structures. In BALB/c mice, adaptation favored nonsynonymous substitutions linked to increased virulence, including a hemagglutinin variant exclusively fixed in female lineages. It also revealed the first demonstration of sex-dependent selection shaping a viral protein interface. In female-adapted viruses, substitutions disrupting a key NS1 dimerization motif converged on a single residue, while in male-adapted viruses, they were dispersed across the same interface. Conversely, adaptation to C57BL/6 resulted in fewer substitutions but promoted defective viral genome formation, leading to reduced cytopathic effect and attenuated virulence. This provides the first *in vivo* evidence that host genotype alone can modulate defective viral genome formation. Our results offer critical insights into host–pathogen interactions and reveal that selective pressures imposed by specific genotype–sex combinations can increase virulence across host genotypes, enabling new epidemiological modeling and disease control strategies.

## INTRODUCTION

In the early stages of viral pandemics, such as the 1918 influenza A virus (IAV) and SARS-CoV-2 pandemics, viruses encounter host populations with minimal pre-existing immunity. In these settings, selection tends to favor mutations that enhance replication and transmission rather than antigenic escape, which typically emerges later as populations acquire adaptive immunity^1–3^. Because innate immune responses differ substantially depending on host genotype and sex, these factors likely shape viral adaptation, influencing virulence, host range, and epidemic trajectory. However, the individual and combined effects of host genotype and sex on viral evolution remain poorly understood.

Host genotype and sex significantly influence immune responses and infection outcomes. For instance, BALB/c and C57BL/6 mice exhibit distinct immune profiles, leading to differences in susceptibility and disease severity^4,5^. Sex also shapes immunity, as females typically mount stronger immune responses than males, leading to faster pathogen clearance but increasing the risk of immunopathology. Conversely, males are often more susceptible to infection but experience less severe immune-mediated damage^6,7^. These host differences likely influence viral evolution, as host resistance and tolerance are thought to select for increased replication and virulence^8^.

Previous studies indicate that pathogen adaptation to resistant host genotypes or hosts with strong immune responses can either increase virulence broadly or cause fitness and virulence trade-offs when infecting novel hosts^8–10^. Similarly, theoretical models suggest that the robust female immune response may select for traits that enhance virulence^11,12^. Dissecting genotype-sex interactions under controlled conditions is essential for understanding how host-imposed selection shapes viral adaptation and virulence. However, direct experimental evidence examining the impact of genotype– sex interactions on pathogen evolution in vertebrate systems is limited^13,14^. The vast diversity of host genotypes in natural populations, combined with the short infection periods of acute viruses, further complicates efforts to understand how host–virus interactions shape pathogen evolution. Experimental evolution offers a powerful approach to address this complexity.

Here, we use experimental evolution to investigate how host genotype and sex shape IAV adaptation in male and female BALB/c and C57BL/6 mice. Utilizing a novel three-dimensional mapping approach, we identify genotype– and sex-specific mutational hotspots and characterize host-dependent evolutionary trajectories. We provide the first evidence of sex-dependent selection patterns and host genotype-specific differences in defective viral genome formation. These results offer new insights into how genotype and sex together drive viral evolution, with broader implications for guiding infection control efforts in genetically diverse populations.

## RESULTS

### The evolution of viral replication phenotypes is host-genotype dependent

To investigate how host genotype and sex affect the evolution of influenza A virus (IAV), we adapted the original H3N2 human isolate A/Hong Kong/1/1968 (HK68) to female and male BALB/cJ and C57BL/6J mice (the adapted viruses are hereafter referred to as BALBF, BALBM, BL6F, and BL6M). After selecting the highest titer viruses from ancestral HK68 infections, we established 7 independent viral lineages per genotype– sex combination and passaged each for 10 rounds, yielding a total of 28 mouse-adapted lineages. This design enabled direct comparisons of evolutionary outcomes across multiple lineages per host group, while controlling for stochastic variation.

First, we wanted to determine whether adaptation to distinct host backgrounds impacts viral fitness, using whole-lung viral load as the measure of viral replication. For each evolved lineage, we measured titers through TCID_50_ assays on lung homogenates collected three days post-infection. Adaptation to BALB/c mice significantly increased titers in both sexes (Fig. 1A), with evolved lineages exhibiting higher lung viral loads than the ancestral HK68 strain (BALBF: U = 306, p < 0.001; BALBM: U = 312, p < 0.001). On the other hand, C57BL/6 adaptation resulted in significantly reduced TCID_50_ values in males (U = 207, p = 0.004) and no significant change in females (U = 169.5, p = 0.156) compared to HK68, indicating a genotype-dependent effect on replication.

**Fig. 1.**
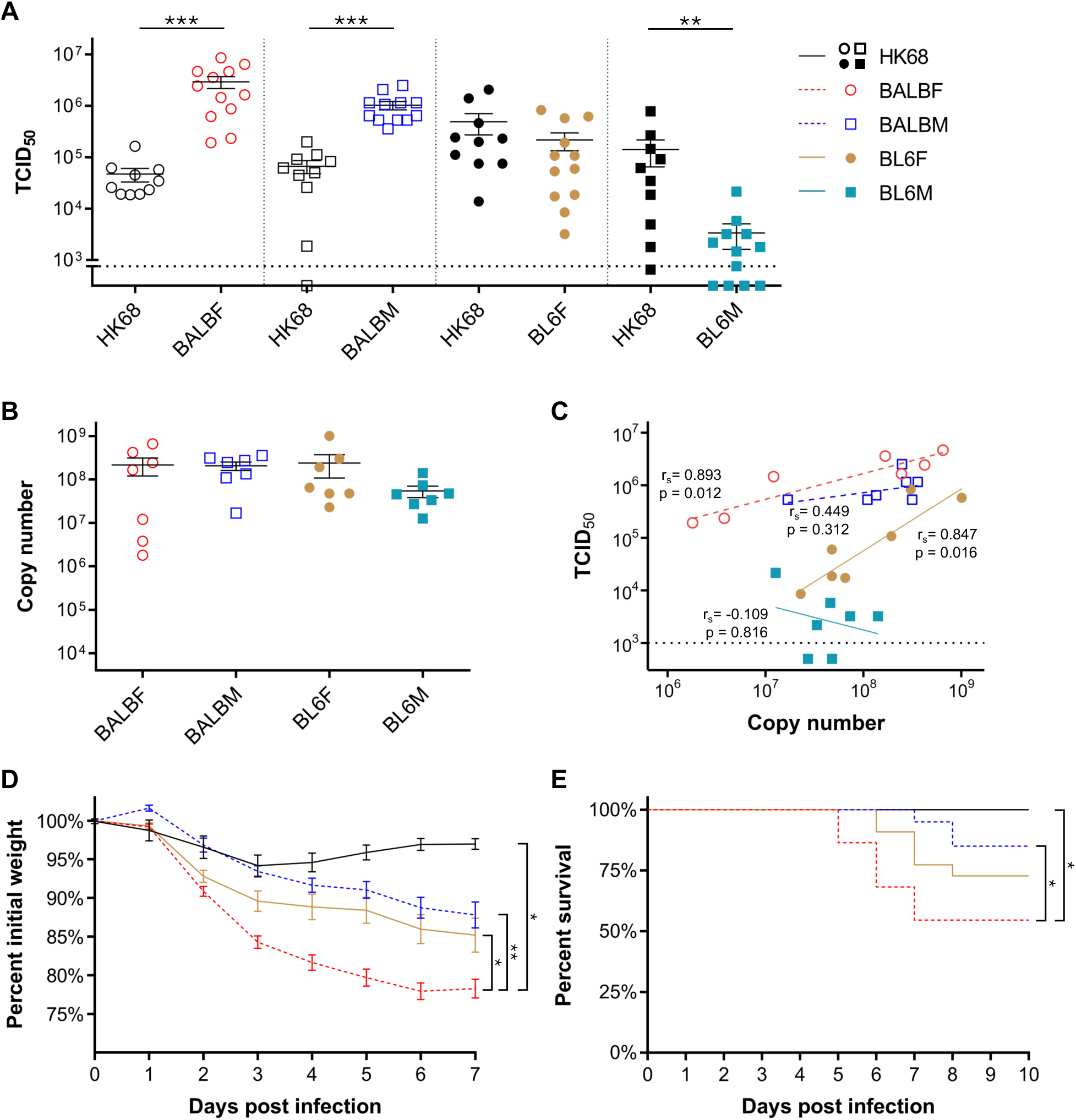
– Titer and virulence measurements. A) Log_10_ TCID_50_ values for viral infections with ancestral HK68 (black, n = 10/host) or mouse-adapted virus (colored, n = 12/host). BALB/c females – open circles; BALB/c males – open squares; C57BL/6 females – solid circles; C57BL/6 males – solid squares. The line at 10^3^ TCID_50_ represents the limit of detection. B) Log_10_ copy number per µL of viral RNA (M segment) (n = 7/host). C) Correlation between log_10_ TCID_50_ and log_10_ vRNA copy number. Spearman’s correlation coefficients (r_s_) were calculated for all lineages of each host-adapted virus (n = 7/host). D) Weight loss (percent of initial weight) and E) survival curves for the ancestral and host-adapted viruses in the respective hosts. Black – Combined ancestral HK68 (n = 29); dashed red line – BALBF infections (n = 22); dashed blue line – BALBM infections (n = 20); solid brown line – BL6F infections (n = 22). Weight loss was modeled until 7 days post infection. Error bars are SEM.

Qualitative RT-PCR was performed after each passage to confirm the presence of virus in lung homogenates, and cycle threshold values for BL6F and BL6M lineages were consistently comparable to those of BALB/c. This indicated that replication was proceeding normally in C57BL/6 mice despite reduced infectivity in cell culture. We therefore hypothesized that adaptation to C57BL/6 abrogated cytopathic effect in MDCK cells, particularly in male-adapted virus. To test this, we performed standardized RT-qPCR on round 10 (R10) samples from each evolved lineage to examine the relationship between viral RNA copy numbers and titers.

Copy numbers were not significantly different among host-adapted viruses (Kruskal– Wallis statistic = 3.7617, p = 0.288), confirming normal replication (Fig. 1B). Furthermore, there was a strong positive correlation between TCID_50_ and copy number in BALBF (r_s_ = 0.893, p = 0.012) and BL6F (r_s_ = 0.847, p = 0.016) lineages, and BL6F titers were roughly an order of magnitude lower for a given copy number (Fig. 1C). By contrast, BALBM lineages showed a similar but statistically non-significant relationship (r_s_ = 0.449, p = 0.312), while BL6M TCID_50_ values were close to or below the limit of detection (10^3^) and there was no correlation between TCID_50_ and copy number in these lineages (r_s_ = –0.109, p = 0.816). The overall similarity in viral genome copy numbers across hosts, together with the significantly reduced titers in C57BL/6-adapted lineages, show that adaptation to different host genotypes and sexes can strongly affect cytopathic phenotypes, influencing traditional viral quantification methods and potentially virulence.

### Host genetic background and sex alter virulence evolution trajectories

We next evaluated whether adaptation to different hosts increased virulence by measuring weight loss and mortality. Because host-associated factors in lung homogenates confounded infection outcomes (data not shown), we first purified each R10 lineage by an overnight cell culture passage (MOI = 0.1), a process that largely preserved viral variants (r^2^ = 0.953, p < 0.001; Fig. S1A) and improved sequence quality (Fig. S1B). After titering, dose-standardized infections were performed in BALBF, BALBM, and BL6F mice. BL6M lineages could not be standardized by TCID_50_ and were therefore excluded from these assays.

Linear mixed models (LMMs) revealed that host-adapted viruses caused more weight loss than the ancestral HK68 in their respective hosts (Fig. 1D). This effect was most pronounced in BALBF, where daily weight loss increased 5.2-fold relative to HK68 (t = –2.868, p = 0.013). BALBM and BL6F lineages induced 5.0– and 3.3-fold greater weight loss, respectively, but these were not statistically significant (p > 0.05), reflecting the high variance in host response. Comparisons between adapted viruses also showed that BALBF was the most virulent overall, causing nearly twice the weight loss of BALBM (t = –3.167, p = 0.005) and BL6F (t = –2.839, p = 0.011), whereas BALBM and BL6F were comparable (t = 0.344, p = 0.735).

Host adaptation also increased mortality (Fig. 1E): BALBF killed 45% of infected mice (log-rank test: χ² = 5.975, p = 0.015), while BALBM and BL6F killed 15% and 27%, respectively (p > 0.05). Because HK68 caused no mortality, we used log-rank tests for comparisons with ancestral virus, and Cox proportional hazards (CPH) between adapted lineages. CPH analysis showed that BALBF induced significantly higher mortality than BALBM (HR = 4.2437, p = 0.028) but not BL6F (HR = 2.113, p = 0.148). These findings indicate that while adaptation increased virulence in all examined hosts, the magnitude of these effects is host-dependent, underscoring the critical role of host genotype and sex in shaping virulence outcomes.

Our experimental design incorporated different pooling treatments to evaluate whether multiple infections during and after adaptation influenced viral fitness and virulence relative to single infections (Fig. S2A). While pooling significantly increased viral titers and mortality (Fig. S2B, C), outcomes varied by host (Fig. S2D–M) and their evolutionary effects were minor compared to those of host genotype and sex; thus, they are not discussed further.

### A rolling sphere model reveals selection hotspots in three-dimensional space

Given that adaptation to different host genotypes and sexes resulted in distinct phenotypes for cytopathic effect and virulence, we asked whether these phenotypic differences arose from underlying genetic variation. To investigate this, we conducted whole genome sequencing of the ancestral HK68 and the 28 independent mouse-adapted R10 viral lineages. Of note, C57BL/6-adapted lineages had higher sequencing coverage than BALB/c lineages, whereas BALB/c samples showed marked degradation, likely due to elevated host RNase activity.

Genomic analysis revealed a broad array of mutations, with non-synonymous variants appearing at various frequencies across lineages (Fig. S3 and Table S1). Several substitutions reached high frequency or even fixation in multiple lineages, suggesting positive selection. While shared variants were readily identifiable, we also encountered instances where high-frequency variants were either tightly clustered or dispersed across the nucleotide sequence. This observation indicated potential “hotspots” arising from broad selective pressures on protein interfaces or entire domains, rather than single substitutions. An initial rolling-window approach to detect genomic regions of elevated non-synonymous variant frequency failed to capture mutations that are distant in linear sequence but in proximity within the three-dimensional architecture of quaternary protein structures.

To overcome this, we devised a “rolling sphere” model that iteratively navigates the protein chains, identifying significant increases in cumulative variant frequency within a defined radius around each residue (<u>Video S1</u>). We applied the model to the entire dataset and separately to host-specific lineages, distinguishing general mouse adaptations from those unique to particular genotypes or sexes. Two sphere radii were used sequentially: a 6 Å radius to detect single amino acid mutations and directly interacting neighbors, followed by a 12 Å radius to encompass broader interfaces and potential allosteric effects. This strategy accommodates interactions within homomeric or heteromeric oligomers and addresses the spatial complexity of viral proteins, enhancing our ability to identify host-specific evolutionary pressures.

### Host genotype and sex determine the emergence of virulence-associated variants

Using the rolling sphere model, we identified multiple residues and hotspots under positive selection, demonstrating its ability to capture adaptive patterns at different scales. Table S2 highlights significant hotspots (99.9th quantile), including the average variant frequency within each host group and known impacts on replication or virulence. To visualize their spatial distribution and potential functional effects, we also mapped mean variant frequencies for each host onto three-dimensional protein structures.

At the residue scale, the rolling sphere model revealed several host-specific adaptations associated with increased virulence and replication. In the hemagglutinin (Fig. 2 – HA), two previously documented mutations within the receptor-binding pocket—G218W and G218E—were present at high frequency almost exclusively in BALB/c females, reaching fixation or near-fixation (>90%) in 5 of 7 BALBF lineages. Although not previously reported as sex-specific, these mutations have only been observed in studies using female BALB/c or CD-1 mice^15–18^, emphasizing the sex– and genotype-specific selective pressures driving their emergence and the importance of including both sexes in pathogen evolution research. Functionally, these mutations have been shown to elevate fusion pH^19^, likely enabling viral fusion earlier during endosome maturation. In contrast, N165D—an HA mutation that removes an N-glycosylation site and is linked to increased virulence^20,21^—was found at high frequency across all other hosts but was only detected at a very low frequency (1.2%) in the single BALBF lineage lacking a G218 mutation, suggesting these mutations may be mutually exclusive.

**Fig. 2.**
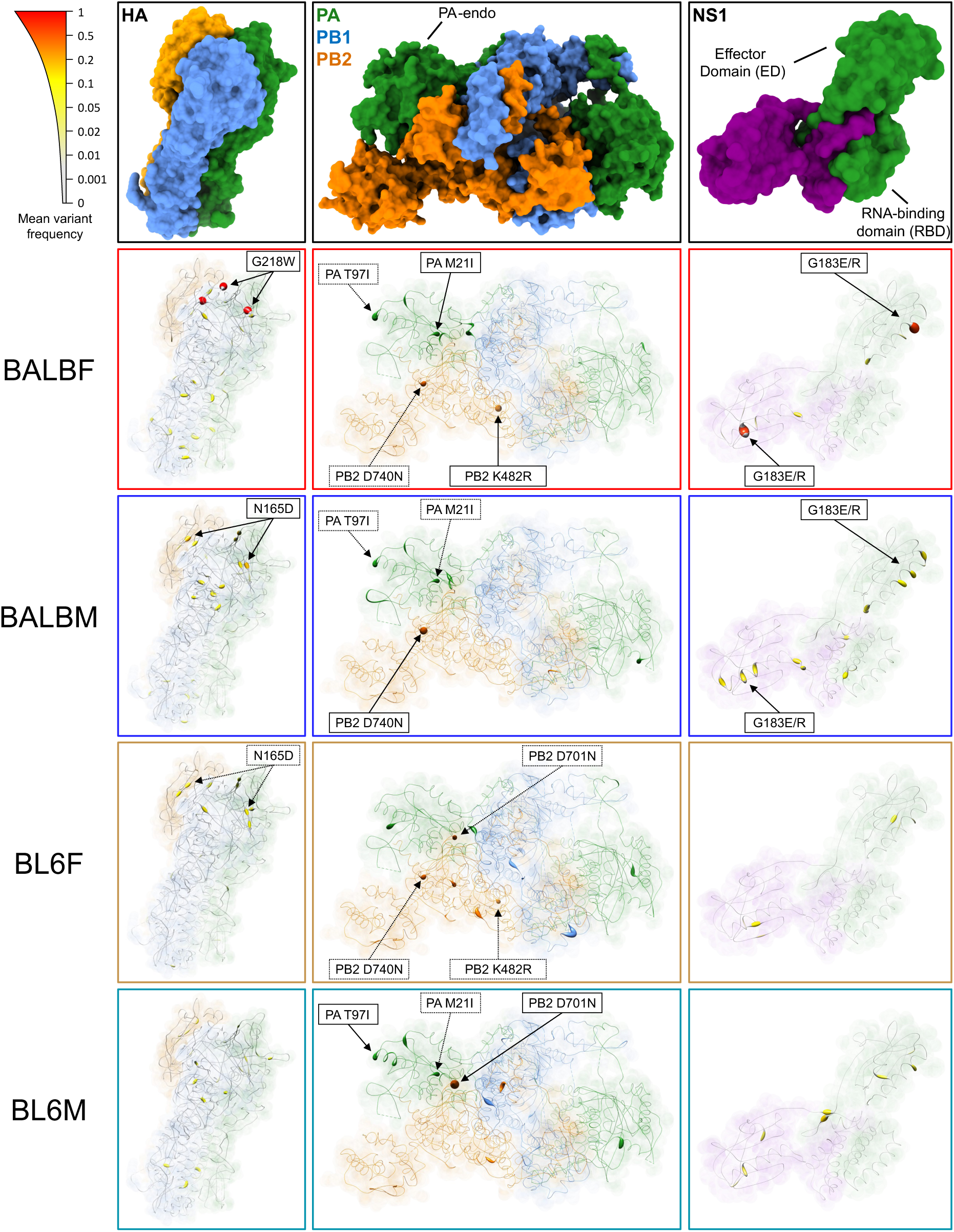
– Three-dimensional protein models for HA (PDB: 7QA4), polymerase (PDB: 6QNW) and NS1 (PDB: 4OPH) adapted to each host. The mean variant frequency across seven independent lineages is represented by the thickness (HA, NS1, polymerase) and color intensity (HA, NS1) of the simplified amino acid chains. Hotspot nexuses are marked with solid lines when statistically significant in a host and dotted lines when present at high frequency in at least one lineage but not statistically significant.

In the polymerase basic 2 (Fig. 2 – PB2), which plays a key role in viral RNA synthesis and host adaptation, we found three positively selected mutations—K482R, D701N, and D740N—all known to enhance polymerase activity or virulence in mammals^22,23^. K482R is localized within the PB2 midlink domain and is involved in nuclear-cytoplasmic shuttling of the polymerase complex. This variant emerged solely in female mice of both genotypes, suggesting a sex-specific adaptation. Meanwhile, D701N and D740N showed clear genotype biases: D701N became fixed or nearly fixed in four C57BL/6-adapted lineages (three male, one female), whereas D740N appeared predominantly in BALB/c (six BALB/c lineages versus one BL6F).

Interestingly, we identified a synonymous M2 mutation of unknown function, Y52Y (TAT→TAC), that although present at 8.4% frequency in HK68, rose to fixation or near-fixation in 23 out of 28 viral lineages across all host types. Neuraminidase (NA), nucleoprotein (NP), matrix protein 1 (M1), and matrix protein 2 (M2) did not show significant nonsynonymous positive selection signals, and the smaller proteins (PA-X, NEP, PB1-F2) lacked suitable Protein Data Bank (PDB) structures and were excluded from the rolling sphere analysis or visualizations.

These findings demonstrate the rolling sphere model’s capacity to detect critical sites under positive selection across distinct host genotypes and sexes. Variants linked to high fitness and virulence arose in specific genotype–sex combinations, indicating that unique host backgrounds can strongly shape the emergence of known mammalian adaptations within a population.

### Spatial analysis provides insights into host-dependent selection at multiple scales

Using two rolling sphere radii (6 Å and 12 Å) further revealed how structural and functional contexts shape viral adaptation. At the interface scale—referring to residue clusters that lie within or near protein–protein interaction surfaces—our model identified a hotspot in the NS1 protein’s effector domain (ED) (Fig. 2 – NS1), which plays key roles in immune evasion and host gene regulation. This hotspot specifically targeted the glycine dimerization motif crucial for ED-ED binding^24^. The site G183 was a significant hotspot at 6 Å in BALBF, harboring G183E/R variants that disrupt a key glycine residue. However, when analyzed with the 12 Å radius, the same interface was also highlighted as a hotspot in BALBM, as distinct charged-residue substitutions were dispersed across the helix-helix interface, in contrast with BALBF lineages, whereas selection in BALBF lineages favored mutations exclusively at G183.

The detection of mutations putatively disrupting NS1 ED-ED dimerization in BALB/c-adapted viruses reflect genotype-specific selection in a domain critical for antagonizing the OAS/RNase L pathway, among other functions^25^. By altering a glycine-based interface with charged residues, these mutations likely impair dimerization and elevate RNase activity, contributing to the observed RNA degradation in BALB/c samples. More strikingly, these genotype-specific mutations were further shaped by host sex: female-adapted lineages had mutations at a single site, while male-adapted lineages exhibited a more dispersed mutation pattern across the interface. To our knowledge, this is the first documented instance of a sex-dependent spatial mutation distribution within a viral protein domain, implying that male and female hosts can exert distinct selective pressures on protein–protein interfaces and, ultimately, on virulence-associated traits.

At the domain level, the rolling sphere model also identified significant hotspots in the Polymerase Acidic (Fig. 2 – PA) endonuclease domain (PA-endo), responsible for cleaving host mRNAs for “cap-snatching”. Two high-frequency variants, M21I and T97I, were found across multiple hosts, alongside numerous lower-frequency substitutions, designating this region as a major hotspot at the 12 Å radius in the combined dataset and specifically in BALB/c-adapted viruses. The high mutational tolerance of PA-endo suggests it may be a suboptimal antiviral target; indeed, we detected three low-frequency variants at residue I38 (I38M/T/V), two of which confer resistance to baloxavir^26^. Such findings underscore the utility of three-dimensional variant scanning for identifying mutable protein sites and guiding targeted drug design.

Our results revealed host-specific selective pressures at various structural scales, thus, we next examined broader evolutionary trajectories using principal component analysis (PCA) on the cumulative variant frequencies within the 6 Å and 12 Å radii centered at each amino acid site of each evolved lineage, and included the ancestral HK68 as a baseline reference (Fig. 3A, B). The PCA revealed clear genotype-specific clustering, with BALB/c-adapted lineages diverging further from HK68 than C57BL/6-adapted lineages. Moreover, BALBF and BALBM lineages exhibited pronounced separation at 6 Å but converged at 12 Å, likely because discrete NS1 ED mutations appear as individual variants on the smaller scale yet coalesce into a single interface-level hotspot at a larger radius. By contrast, C57BL/6 lineages remained comparatively closer to both HK68 and each other, suggesting more limited positive selection. These genotype-specific differences were further supported by additional metrics of viral evolution: BALB/c-adapted lineages accumulated significantly more nonsynonymous polymorphisms (U = 163, p = 0.0029) and exhibited greater frequency-weighted divergence (U = 149, p = 0.0186) from HK68 than C57BL/6 lineages (Fig.3C, D).

**Fig. 3.**
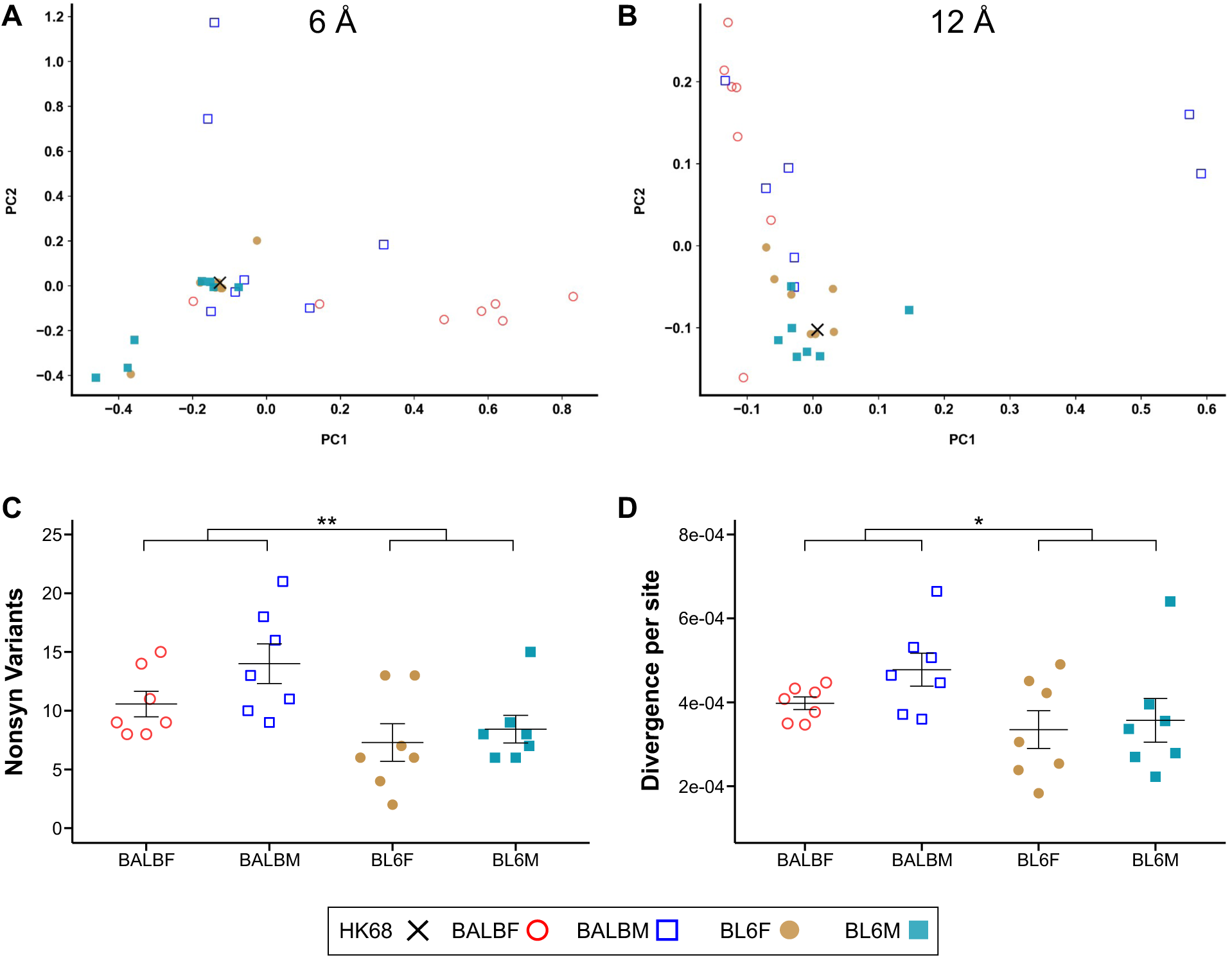
– A, B) PCAs for the sum of cumulative frequencies at each site for 6 Å (A) and 12 Å (B) for each host group. Ancestral HK68 is marked as a black X. C) Sum of nonsynonymous variants above 5% frequency for each viral lineage. D) Frequency-weighted genetic divergence per site for each evolved sample compared to HK68. n = 7 per host type for all analyses. Error bars are SEM.

These results highlight that analyzing variation across structural scales resolves selection patterns that may be missed at either level alone. Residue-scale analyses can overlook broader structural convergence, while interface-scale approaches may miss host-specific selection at individual sites. By integrating spatial and functional information, the rolling sphere model clarifies how genotype and sex shape viral adaptation and drive the emergence of virulence-associated variants.

### Host genotype and sex influence defective viral genome formation

As C57BL/6-adapted lineages lacked shared substitutions or hotspots that could explain their distinctive loss of cytopathic effect, we investigated alternative mechanisms underlying this phenotype. A known contributor to reduced tissue culture infectivity is the accumulation of defective viral genomes (DVGs) that can be packaged into virions, forming defective interfering particles. Previously considered laboratory artifacts^27–29^, DVGs are now recognized as naturally occurring elements that interfere with replication, modulate immune responses, and influence viral persistence and virulence *in vivo*^30,31^. Recent studies confirm that DVGs are present in human influenza infections, highlighting their relevance to host immunity and clinical outcomes^32–34^.

To assess whether DVGs were responsible for reduced cytopathic effect, we screened for deletions (≥3 nucleotides) across viral genomes from R10 lung homogenates.

Deletions primarily occurred near the terminal regions of polymerase segments (PB1, PB2, PA), consistent with known influenza DVGs^35^ (Fig. 4A–D). Host genotype significantly influenced DVG formation, with C57BL/6 viruses having more numerous (U = 414.5, p < 0.001) and longer deletions (U = 1156.5, p = 0.021) compared to BALB/c viruses. Additionally, male-adapted viruses had longer deletions than female-adapted viruses (U = 1524, p = 0.029), aligning with the lower TCID_50_ values observed in male lineages, particularly BL6M. These results, detailed in Table S3, reveal the substantial influence of host genotype and sex on DVG formation.

**Fig. 4.**
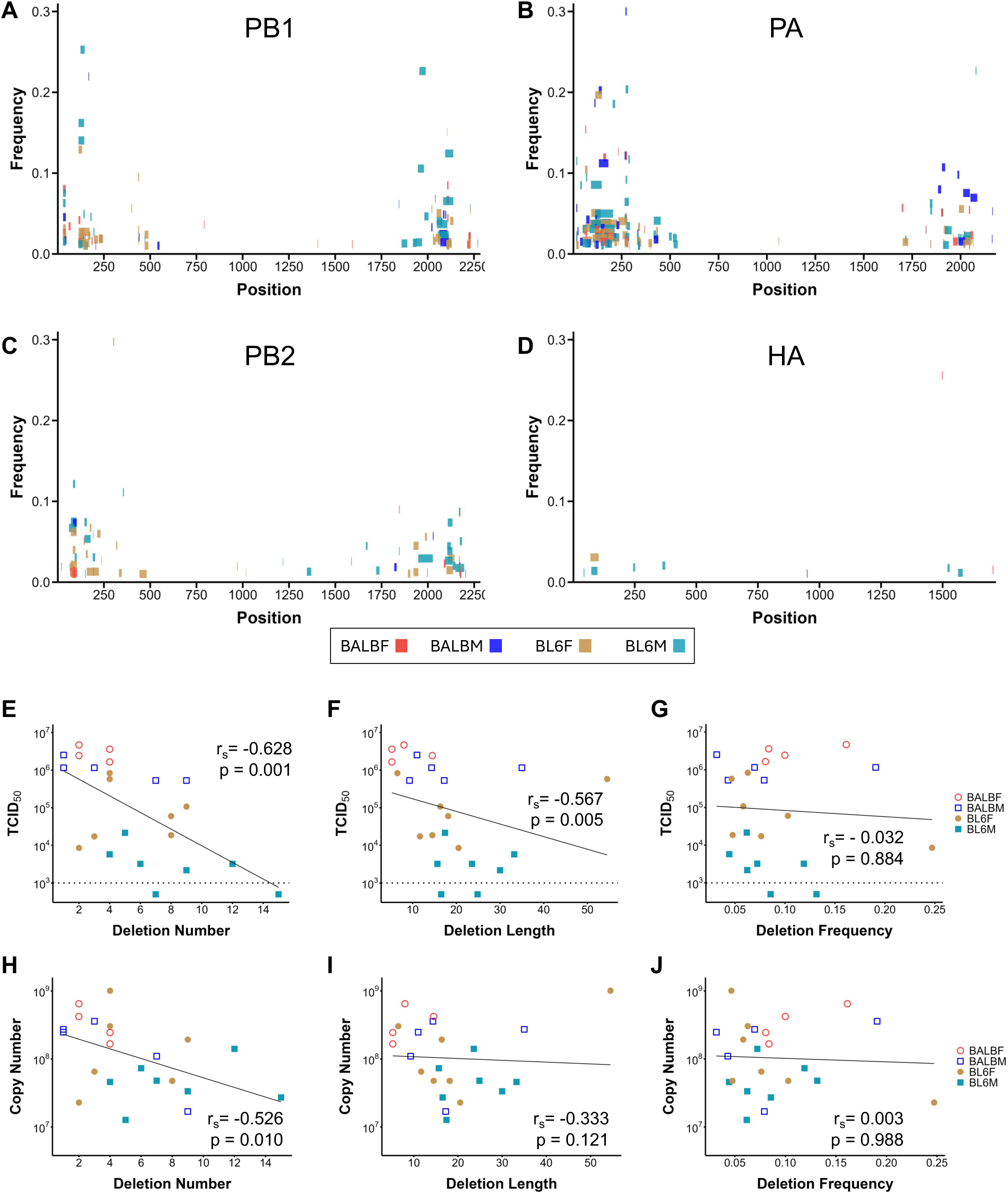
– Genomic deletion measures by host. A-D) Genomic position (x-axis), Frequency (y-axis) and length (width) of deletion variants for the polymerase segments and hemagglutinin. Lineages within each host are combined. E-J) Spearman’s correlations between TCID_50_ and deletion number (E), deletion length (F) and deletion frequency (G), and correlations between genome copy number and deletion number (H), deletion length (I) and deletion frequency (J). Samples with mean polymerase coverage <100 reads are excluded. n = 7 per host type.

TCID_50_ negatively correlated with both deletion number (r_s_ = –0.628, p < 0.001) and length (r_s_ = –0.567, p = 0.005) (Fig. 4E–G), confirming their impact on infectivity. Moreover, genome copy number negatively correlated with deletion number (r_s_ = –0.526, p = 0.010) (Fig. 4H–J), suggesting that DVGs may have also impaired replication. These findings provide the first evidence for host-induced differences in DVG formation *in vivo*, revealing that viral adaptation can not only occur through selection of genetic variants but also by exploiting host-specific immune mechanisms that balance persistence and virulence.

### Viral adaptation to specific host genotypes increases virulence in all hosts

Having established that adaptation to distinct host genotypes generates viral populations that are not only genetically divergent but also exhibit varying levels of genetic diversity and DVG content, we next evaluated virulence in the original (familiar versus novel (unfamiliar) host environments. To this end we infected female C57BL/6 mice with the BALBF lineages, and female BALB/c mice with the BL6F lineages, to assess weight loss and mortality outcomes compared to familiar virus infections.

BALBF virus induced significantly greater weight loss (LMM; t = 2.388, p = 0.049) and higher mortality (CPH; HR = 3.198, p = 0.001) compared to BL6F virus, independent of the recipient host genotype (Fig. 5A, B). While BALBF virus caused similar mortality in both host environments (log-rank; χ² = 1.390, p = 0.238), BL6F virus exhibited significantly reduced mortality when introduced into BALB/c mice (log-rank; χ² = 4.364, p = 0.037).This suggests that while C57BL/6-adapted virus showed decreased virulence in a novel host, the BALB/c-adapted virus retained high virulence across host genotypes.

**Fig. 5.**
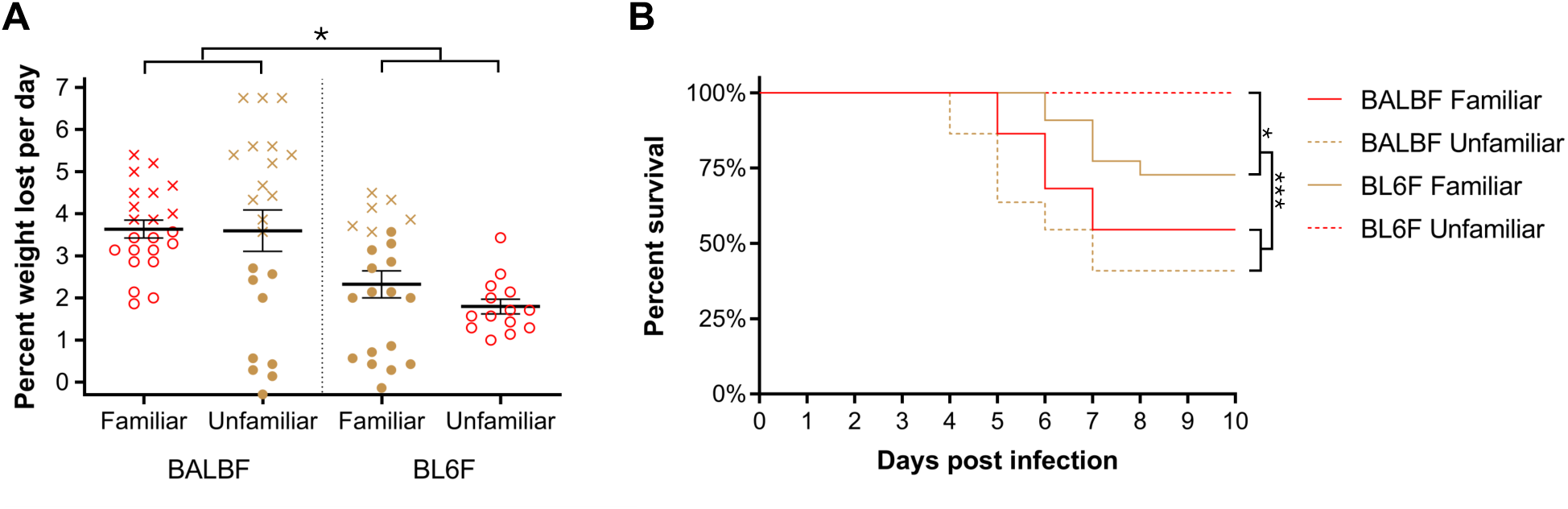
– A) Weight loss (mean percent weight lost per day) and B) survival curves for familiar (solid) and unfamiliar (dashed) host infections with BALBF and BL6F virus. BALB/c female mice – open red circles/red lines; C57BL/6 female mice – solid brown circles/brown lines. Mice that died are represented as crosses. BALBF familiar (n = 22); BALBF unfamiliar (n = 22); BL6F familiar (n = 22); BL6F unfamiliar (n = 14). Error bars are SEM.

Overall, C57BL/6 female mice experienced slightly higher mortality than BALB/c females for both familiar and unfamiliar viruses, although this difference was not statistically significant (CPH; HR = 1.407, p = 0.159). This outcome is consistent with the heightened variability of C57BL/6 immune responses, which can either mitigate infection effectively or exacerbate immunopathology and increase mortality. This bimodal response, evident in Fig. 5A, increased variance and reduced statistical power in some comparisons despite clear differences in mean weight loss. In contrast, BALB/c responses were more uniform, resulting in lower variance and more consistent infection outcomes. These data highlight the complex interplay between host-specific immune dynamics and viral virulence across familiar and novel environments.

Collectively, these results demonstrate that while variation in individual host immune responses influences whether viral infection is mitigated early, a virus’s evolutionary history—shaped by prior host immune pressures—is the primary determinant of its overall virulence. Certain host genotypes select for high-virulence variants, exemplified by the BALB/c female-specific G218W/E mutations, and this elevated virulence can persist even in genetically distinct hosts, regardless of the specific host environment.

## DISCUSSION

Using experimental evolution, we previously demonstrated that adapting Friend virus to distinct host genotypes or MHC backgrounds resulted in fitness and virulence trade-offs when infecting novel hosts^9,10,36,37^. Extending this work, we investigated how host genotype and sex interact to shape IAV evolution. Our study revealed two distinct evolutionary trajectories driven by host genotype. Ancestral HK68 virus initially had lower fitness and virulence in BALB/c compared to C57BL/6 mice, but adaptation to BALB/c selected variants associated with increased virulence across both host genotypes, supporting predictions that resistant hosts may drive selection for increased virulence. Conversely, C57BL/6 adaptation primarily resulted in reduced cytopathic effect. While these phenotypic differences were evident, understanding the underlying genetic basis required moving beyond traditional linear genomic analyses.

Mapping mutations to structural models is common, however, most existing tools rely on visual identification of mutational hotspots within individual proteins and lack genome-wide, automated hotspot detection^38–40^. Our rolling sphere model enables automated identification of selection hotspots across entire viral genomes, given sufficient structural data. Applied to our dataset, the model revealed genotype– and sex-specific selective pressures across viral proteins and domains. Some mutations, such as PB2 K482R, D701N, and D740N, displayed broad genotype-or sex-associated patterns across hosts. Others reflected strong genotype–sex interactions, most notably in BALB/c-adapted viruses. In hemagglutinin, receptor-binding pocket mutations (G218W/E) were nearly exclusive to BALB/c female lineages, suggesting selection for accelerated viral entry in this host. In NS1, mutations putatively disrupting ED–ED dimerization also occurred in BALB/c-adapted viruses but exhibited a sex-dependent distribution: mutations in female-adapted lineages were concentrated at a single residue (G183) but were dispersed across the interface in male viruses, indicating convergent outcomes shaped by slightly distinct selective pressures. This represents, to our knowledge, the first report of a sex-dependent spatial selection bias at a host–pathogen interface.

Although viral mutation rates are high and every possible mutation likely occurs multiple times during an infection, purifying selection, genetic drift and within– and between-host bottlenecks typically limit nonsynonymous mutation transmission^41,42^. Therefore, infections in hosts that strongly select for virulence-associated variants, such as BALB/c females with HA G218W/E, may be crucial for these variants to reach transmissible frequencies. Understanding intra-host competition among viral variants will be key to determining whether such host-favored substitutions enhance viral fitness across host genotypes and whether they persist in genetically diverse populations. If so, this would imply that certain hosts can act as evolutionary accelerators, triggering punctuated evolutionary bursts with broad epidemiological consequences.

Beyond the selection of single-nucleotide variants, host genotype—and to a lesser extent sex—significantly influenced defective viral genome (DVG) formation. While recent cell culture studies have suggested host factors can modulate DVG production^43,44^, this study offers the first *in vivo* evidence of genotype-specific differences in DVG accumulation. DVG-rich cells stimulate antiviral immunity while inhibiting apoptosis, potentially reducing virulence but extending infection duration^45^. C57BL/6-adapted lineages accumulated more frequent and longer deletions, consistent with increased DVG content, likely contributing to reduced cytopathic effect and virulence. Moreover, enhanced DVG formation may limit the effective viral population size, promoting purifying selection and constraining genetic diversity and adaptability. This potentially contributes to the reduced divergence from ancestral HK68 observed in C57BL/6 viruses. These findings highlight a host-driven mechanism of viral attenuation with important implications for persistence, evolution, and disease outcomes.

In conclusion, our study demonstrates that host genotype and sex profoundly shape influenza A virus evolution through mechanisms including sex-biased selection at protein interfaces and host-specific differences in DVG formation. These findings reveal how host-driven selective pressures promote the emergence and maintenance of virulence-associated variants. Identifying the specific host mechanisms involved will be key to predicting viral evolution, improving epidemiological models, and developing control strategies that account for host-driven selection.

## Methods

### Mice

Infections were conducted on age-matched 7–24-week-old C57BL/6J and BALB/cJ mice of both sexes, sourced from The Jackson Laboratory or bred in-house from Jackson Laboratory ancestry. Mice were acclimated for at least 3 days, provided ad libitum access to food and water, and singly housed for at least 24 hours prior to infection. A total of 686 mice were used. All procedures adhered to University of Utah IACUC guidelines.

### Viral rescue, propagation, purification and quantification

Influenza strain A/Hong Kong/1/1968 (HK68) was provided by Dr. Earl Brown (University of Ottawa) in the form of 8 plasmids, each containing an influenza segment. Viral rescue was performed by Matthew Angel and Jonathan Yewdell (NIH/NIAID, Bethesda, MD) following a previously described protocol with minor modifications^46^. Briefly, Lipofectamine 2000 (Invitrogen) and DNA (1 µL:1 µg) were incubated for 30 min at RT. Opti-MEM (Gibco-BRL) was added to 1 mL total volume and applied to MDCK/293T co-culture. After 6 h, the medium was replaced with fresh Opti-MEM. At 24 hours post-transfection, 2 mL of Opti-MEM containing TPCK-trypsin (1 µg/ml) was added. At 3 days post-transfection, supernatant was harvested and inoculated into the allantoic cavity of 10-day-old SPF embryonated chicken eggs. At 48 hours post-inoculation, virus-containing allantoic fluid was harvested and stored at –80 ⁰C.

Viral titers were measured using standard TCID_50_ assays in MDCK cells and calculated using the Reed-Muench method^47,48^. For intermediate passages, presence of virus in samples was determined by extracting RNA (Quick-RNA Miniprep, Zymo Research) and performing quantitative reverse transcription polymerase chain reaction (RT-qPCR) with the Verso 1-step RT-qPCR Kit (Thermo Fisher) using primers specific for the M segment of IAV^49^.

Evolved lineages were purified using a custom cell-culture protocol prior to testing. Briefly, 6-well plates with confluent MDCK cells were inoculated with 400 µL of viral growth medium containing the post-passage lung homogenates (MOI = 0.1). After a 60 minute adsorption, lung-derived medium was washed off, replaced with new viral growth medium and incubated overnight at 37 ⁰C. After this incubation, medium was aspirated into a 2mL microtube, centrifuged for 5 minutes at 300 × g, 4 ⁰C, and the supernatant collected, mixed, aliquoted, and stored at –80 ⁰C.

MDCK cell maintenance medium was comprised of DMEM with high glucose, L-glutamine and sodium pyruvate (Genesee Scientific); 10% FBS, heat-inactivated (Gibco); 1% Penicillin-Streptomycin (Sigma-Aldrich, St. Louis, MO).

Viral growth medium was DMEM as above; 2.5% BSA (Fisher Scientific); 2.5% HEPES (Fisher Scientific); 1% Pen-Strep; 2 µg/mL TPCK-trypsin.

### Viral passage and collection

For serial passages, ten mice of each host type were infected with ancestral HK68 and the four viruses with the highest lung titers in each group were selected and designated as round 1 (R1) A, B, C, and D. These four initial viruses served as the starting point for seven independent viral lineages in each host group. Viruses were passaged for 10 rounds of infection using three different strategies: single-host passages (1X), where viruses from R1 A, B, C, and D were passaged independently for 10 rounds; two-host pooled passages (2X), where viruses from R1 A and B were pooled to create lineage AB, and viruses from R1 C and D were pooled to create lineage CD; and four-host pooled passages (4X), in which viruses from R1 A, B, C, and D were all pooled to create lineage ABCD. This was done separately for each host, resulting in a total of 28 independent viral lineages.

Passages followed a modified standard protocol^50^ with 20 µL intranasal inoculations under isoflurane anesthesia. Each passage was conducted using 20 µL intranasal inoculations in mice under isoflurane anesthesia. After 3 days mice were sacrificed and lungs extracted into a 2 mL microtube, homogenized with a rotor-stator homogenizer (Dremel) and brought up to a volume of 1 mL using PBS. Homogenates were centrifuged and supernatant collected, mixed, aliquoted, and stored at –80 ⁰C.

To measure virulence, body weights were recorded daily (±2h). Mice reaching ≤75% of initial weight were euthanized per humane endpoints. For test phases, weights were recorded for 10 days or until mice reached the weight threshold or showed prominent signs of distress.

All passages and test phases had animals balanced by litter, age and origin (ordered from The Jackson Laboratory or bred in our facility), to avoid any potential bias across treatments. All test phase mice received standardized doses of 4000 TCID_50_ per inoculate.

### Viral titer and virulence statistics

Viral titers were assessed using two distinct nonparametric approaches to account for non-normality and unequal variance that could not be addressed with transformation. For each host group, a Wilcoxon-Mann-Whitney test compared combined passage treatment titers (1X, 2X, 4X) against stock virus. Passage treatments were compared by Kruskal-Wallis tests, followed by Dunn’s multiple comparison test (dunn.test). For evaluation of association between viral copy number and TCID_50_, we employed Spearman’s rank correlation to accommodate non-linear relationships without assuming data normality.

Test-phase virulence was assessed using linear mixed models (LMMs) to predict body weight as a percentage of initial weight. The models included fixed effects for time (day postinfection), passage status (Passaged or Unpassaged), and host group (BALBF, BALBM, BL6F), as well as their interactions. Observations were nested within mouse ID and further nested within viral lineage to reflect the hierarchical structure of the data. Given LMMs’ assumption of linearity, the analysis was limited to the first 7 days post-infection, as recovery in surviving mice leads to restored body weight. Independent analyses were conducted for different pooling treatments (1X, 2X, 4X, 1XP) and for familiar versus unfamiliar infections. For these analyses, host of adaptation was included as a fixed effect where appropriate. All LMMs were fitted using the lme4 package in R, with degrees of freedom estimated via Satterthwaite approximation (lmerTest). Model selection was guided by corrected Akaike’s Information Criterion (AICc) using the MuMIn package.

Differential mortality was assessed using Cox proportional hazards models with similar structures to the body weight analyses. For pairwise comparisons where the Cox proportional hazards model failed to converge due to an absence of mortality, log-rank tests were performed instead. Both tests done using the R survival package.

### Sequencing

Sequencing preparation and read processing followed a previous protocol^41^ with modifications. Briefly, Viral RNA was extracted using the Purelink Pro 96 Viral RNA/DNA Purification Kit (Thermo Fisher) with automated processing on the EpMotion system, followed by elution in 100 µL of RNase-free water. RNA integrity was confirmed via multiplex RT-PCR using SuperScript III One-Step RT-PCR System with Platinum Taq High Fidelity DNA Polymerase (Thermo Fisher), amplifying all eight influenza A virus gene segments. PCR products were purified using Agencourt AMPure XP magnetic beads and verified via agarose gel electrophoresis.

Library preparation was conducted using the Illumina Nextera DNA Flex Library Prep Kit, followed by size selection and quantitation. Indexed libraries were pooled, and qRT-PCR was used to determine viral genome copy number prior to sequencing.

For read processing, adapters were removed using cutadapt, and reads were aligned to the HK68 reference genome with bowtie2. Duplicates were marked and removed using Picard and samtools, and variants were called using deepSNV. Filtering criteria included a p-value < 0.05, Phred quality score >35, mapping quality (MapQ) >30, and read positions between 40 and 200. Samples with low sequencing coverage were excluded from further analysis. All WGS data used in this manuscript can be accessed in BioProject PRJNA1228145.

### Rolling sphere model

Selection hotspots were identified using a three-dimensional rolling window-style approach that calculates the sum of weighted variant frequencies within two defined radii (6 and 12 Å) around each residue in protein structures. The 6 Å radius captured local substitutions and direct residue interactions, while the 12 Å radius identified broader mutational clusters and potential allosteric effects. The model was applied separately to all viral lineages (all hosts combined), genotype groups (BALB/c, BL6), sex groups (males, females), and individual host groups (BALBF, BALBM, BL6F, BL6M) to identify selection patterns at multiple levels.

To determine significance, 20 randomizations were performed, recalculating variant sums at each site for all hosts combined. Sites exceeding the maximum randomized value were considered significant hotspots. Sites containing more than 67% of the summed frequencies within a sphere were designated as the hotspot nexuses and used to distinguish whether selection targeted single amino acids or broader structural regions. From the sites identified using all host data, we conducted post hoc model iterations to identify mutations that were significant at the genotype, sex or individual host level. Those detected at high frequency in both individual host groups were classified as genotype-or sex-specific accordingly, while those present in only one host type were considered host-specific.

### 3D modeling and distance matrices

UCSF Chimera v1.16 and ChimeraX v1.9 were used for 3D visualization of molecular structures. B-factors in the protein data bank (PDB) files were replaced with the mean mutation frequency of all independently evolved lineages in each host group. Distance matrices for the rolling sphere were generated using the PDBparser module from the Bio.PDB library in Python to compute pairwise distances between residue centroids.

The PDBs used for visualization and/or distance matrix generation were: HA – 7QA4 (3D model) and 4WE4 (distance matrix); NS1 – 4OPH; NP – 6J1U; PB1/PB2/PA – 6QNW; NA – 7U4E; M1 – 6Z5J; and M2 – 2RLF.

### Evolutionary metrics

To quantify genetic divergence from HK68, we quantified non-synonymous polymorphisms per sample and also calculated the frequency-weighted divergence for each evolved viral population. Divergence was calculated across all genomic sites using the formula *p*(1 − *q*) + *q*(1 − *p*), where *p* and *q* represent the variant frequency in the evolved and ancestral populations, respectively. Values were summed genome-wide and normalized by the total genome length. To reduce noise from low-frequency mutations, we applied a variant frequency cutoff of 5%, excluding all variants below this threshold from both metrics. Analyses were repeated with and without the cutoff, and results were qualitatively similar, indicating that key patterns of divergence were not driven by low-frequency variants. Host, genotype, and sex effects on both metrics were tested using Kruskal–Wallis or Wilcoxon rank-sum tests as appropriate, with post hoc Dunn’s tests for significant multi-group comparisons.

### Defective viral genome analyses

Genomic deletions were analyzed using conservative cutoffs. Samples with average coverage below 100 reads across the polymerase segments were excluded. Only deletions with a frequency of at least 3% and a minimum length of 3 nucleotides were included in the analysis. The average number of deletions, average deletion length and average deletion frequency per sample were calculated for each host group.

Differences between groups were assessed using a Kruskal-Wallis test, followed by Dunn’s multiple comparison test (dunn.test) for significant pairwise differences. For comparisons involving host genotype and host sex, Wilcoxon-Mann-Whitney tests were performed. Relationships between deletions and TCID_50_ or copy number were assessed using Spearman’s correlations. All plots were generated with R package ggplot2.

## Lead contact

Further information and requests for resources and reagents should be directed to the lead contact, Rodrigo Costa.

## Data and code availability

Raw sequencing data were deposited in the NCBI Sequence Read Archive, BioProject ID [PRJNA1228145] (https://www.ncbi.nlm.nih.gov/bioproject/1228145).

R scripts and data for structural mapping and statistical analyses can be accessed at: https://github.com/RodrigoMamedeCosta/flu-evolution.

Additional information is available from the lead contact upon request.

## Ethics statement

All procedures were approved by the Institutional Animal Care and Use Committee (IACUC) at the University of Utah under protocol no. 17–07021. Experiments were conducted in accordance with the NIH Guide for the Care and Use of Laboratory Animals.

## ACKNOWLEDGEMENTS

The authors thank Dr. Earl Brown (University of Ottawa) for providing the HK68 viral plasmids, Dr. Matthew Angel and Dr. Jonathan Yewdell (NIH/NIAID) for rescuing the virus and Dr. Jody Rosenblatt (King’s College, London) for the MDCK cells. We thank Dr. James Ruff, Dr. Douglas Cornwall and Dr. Joseph Cauceglia for their contributions to the study’s design and statistical analysis.

W.K.P. and F.R.A. were supported by the National Institutes of Health grant R01GM109500. R.M.C. was partly supported by the John H. Weis Memorial Graduate Fellowship. K.C. and K.Z. were supported by the Undergraduate Research Opportunities Program (UROP) at the University of Utah. A.S.L. was supported by a Burroughs Wellcome Fund Investigator in the Pathogenesis of Infectious Diseases award.

## AUTHOR CONTRIBUTIONS

Conceptualization and experimental design: R.M.C., J.S., F.R.A. and W.K.P.; manuscript writing: R.M.C.; editing and revisions: R.M.C., B.S.L., A.S.L., J.S., F.R.A. and W.K.P.; viral passage, purification, processing and data collection: R.M.C., L.A.-A., K.C., K.Z. and J.K.; sequence analysis and statistical analyses: R.M.C., J.S. and F.R.A.; rolling sphere model development and implementation: R.M.C. and F.R.A.; Data visualization and figure preparation: R.M.C. and B.S.L.; Library preparation and viral sequencing: R.M.C, A.L.V., W.J.F., and A.S.L.

## DECLARATION OF INTERESTS

The authors declare no competing interests.

Resources table.

| Resources | Source | Identifier |
| --- | --- | --- |
| C57BL/6J mice, male and female | The Jackson Laboratory | 000664 |
| BALB/cJ mice, male and female | The Jackson Laboratory | 000651 |
| A/Hong Kong/1/1968 (HK68) Influenza Strain | Earl Brown, University of Ottawa<br>Jonathan Yewdell, NIAID | N/A |
| MDCK cells | ATCC | CCL-34 |
| MDCK cells | Jody Rosenblatt, King's College London | N/A |
| 293T cells | ATCC | CRL-11268 |
| Reagents | Source | Identifier |
| Lipofectamine 2000 | Invitrogen | 11668027 |
| Opti-MEM | Gibco-BRL | 31985062 |
| TPCK-trypsin | Worthington | LS003740 |
| Quick-RNA Miniprep Kit | Zymo Research | R1055 |
| Verso 1-step RT-qPCR Kit | Thermo Fisher | AB4101 |
| DMEM | Sigma-Aldrich | D6429 |
| DMEM (high glucose, L-glutamine, sodium pyruvate) | Genesee Scientific | 25-507 |
| FBS (heat-inactivated) | Gibco | 16140071 |
| Penicillin-Streptomycin | Sigma-Aldrich | P4333 |
| BSA | Fisher Scientific | BP1600100 |
| HEPES | Fisher Scientific | BP310 |
| Purelink Pro 96 Viral RNA/DNA Purification Kit | Thermo Fisher | K183001 |
| SuperScript III One-Step RT-PCR System | Thermo Fisher | 12574026 |
| Agencourt AMPure XP beads | Beckman Coulter | A63880 |
| Illumina Nextera DNA Flex Library Prep Kit | Illumina | 20018704 |
| Software and algorithms | Source | Version |
| Cutadapt | PyPI<sup>51</sup> | 5.0 |
| Bowtie2 | SourceForge<sup>52</sup> | 2.5.1 |
| Picard | Broad Institute GitHub<sup>53</sup> | 2.27.4 |
| Samtools | HTSlib GitHub<sup>54</sup> | 1.20 |
| deepSNV | Bioconductor<sup>55</sup> | 1.42.0 |
| PDBparser (Biopython) | Biopython<sup>56</sup> | 1.79 |
| ggplot2 | CRAN<sup>57</sup> | 3.5.1 |
| dunn.test | CRAN<sup>58</sup> | 1.3.6 |
| lme4 | CRAN<sup>59</sup> | 1.1-36 |
| lmerTest | CRAN<sup>60</sup> | 3.1-3 |
| survival | CRAN<sup>61</sup> | 3.8-3 |
| MuMIn | CRAN<sup>62</sup> | 1.47.1 |
| UCSF Chimera | UCSF<sup>63</sup> | 1.16 |
| ChimeraX | UCSF<sup>64</sup> | 1.9 |

**Fig. S1.**
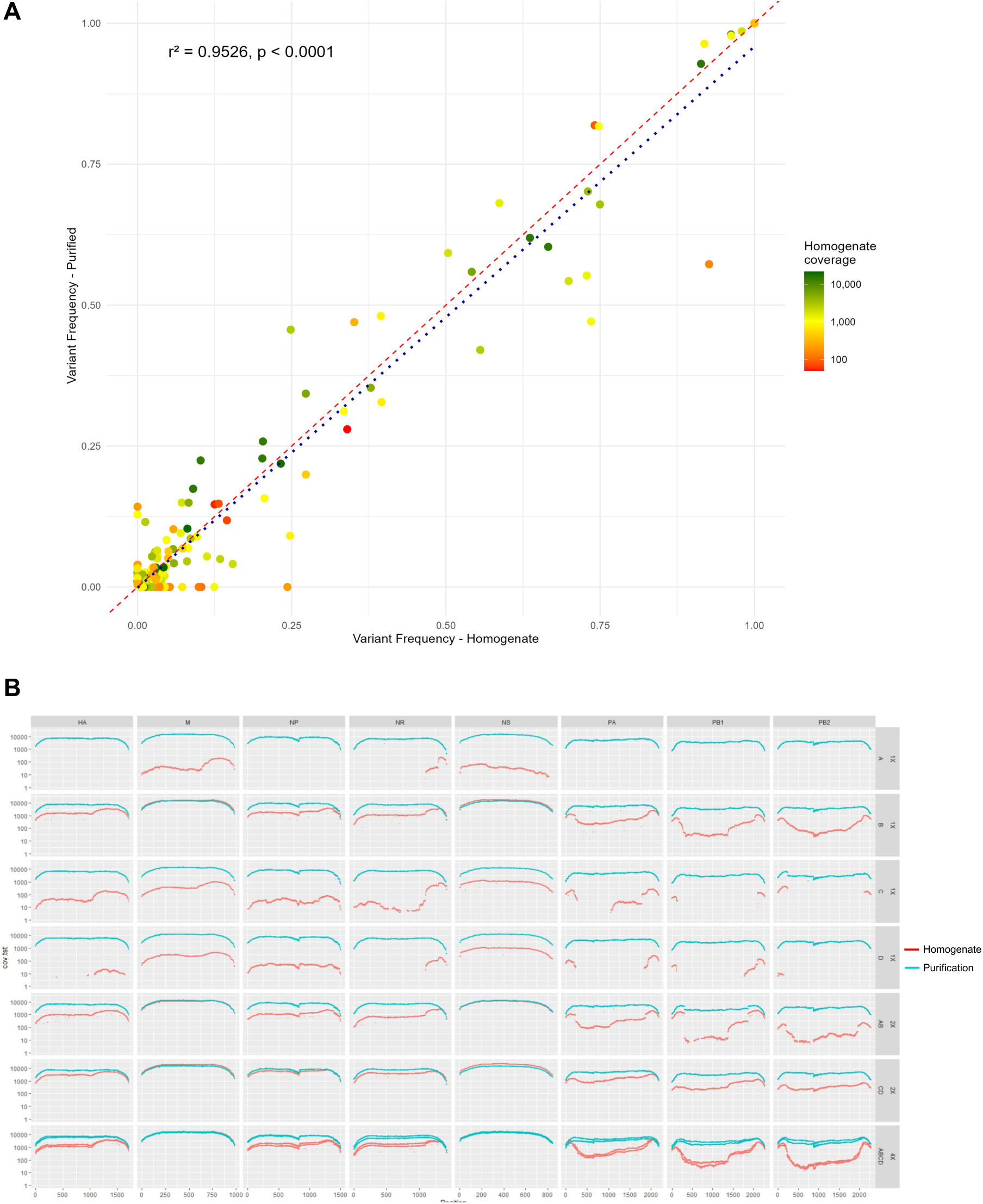
– A) Comparison of variant frequencies in homogenates and purified samples. Each point represents a single variant, with the x-axis showing the frequency in homogenate samples and the y-axis showing the frequency in the respective purified samples. The color of each point indicates the coverage at that variant, (red = low coverage, green = high coverage). The red line represents the line of perfect concordance (y = x), indicating identical frequencies between the two datasets. The blue dotted line is the linear regression line, showing the observed relationship between the two sample sets (r² = 0.9526, p < 0.0001). B) Coverage maps for homogenates (red) and purified (blue) samples. Each column represents a genomic segment, and each row a viral lineage (n = 7).

**Fig. S2.**
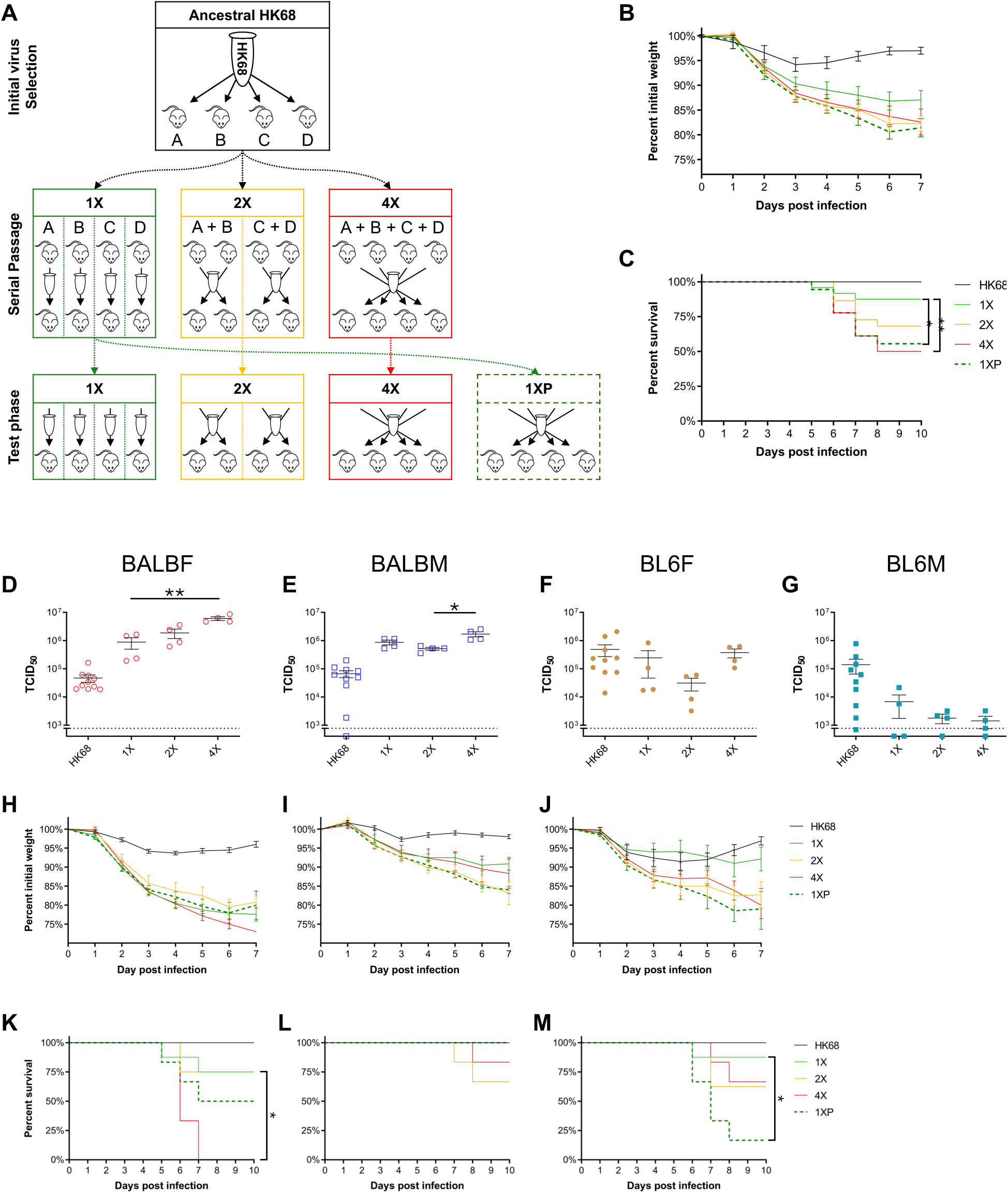
– A) Experimental design. HK68 virus (black) was used to infect ten mice of each host type. The four highest titer viruses in each host were selected to start all passage lineages for that host. B) Combined host data for weight loss and C) survival curves for ancestral (n = 30) and passage/test phase treatments. 1X – single-host passages (green, n = 24); 2X – two-host pooled passages (gold, n = 22); 4X – four-host pooled passage (red, n = 18). 1XP – evolved 1X lineages pooled at the test phase only (dark green, dashed, n = 18). D-G) Titers, H-J) weight loss and K-M) survival for the individual host strains. The respective host is indicated above the different measures. Error bars are SEM.

**Fig. S3.**
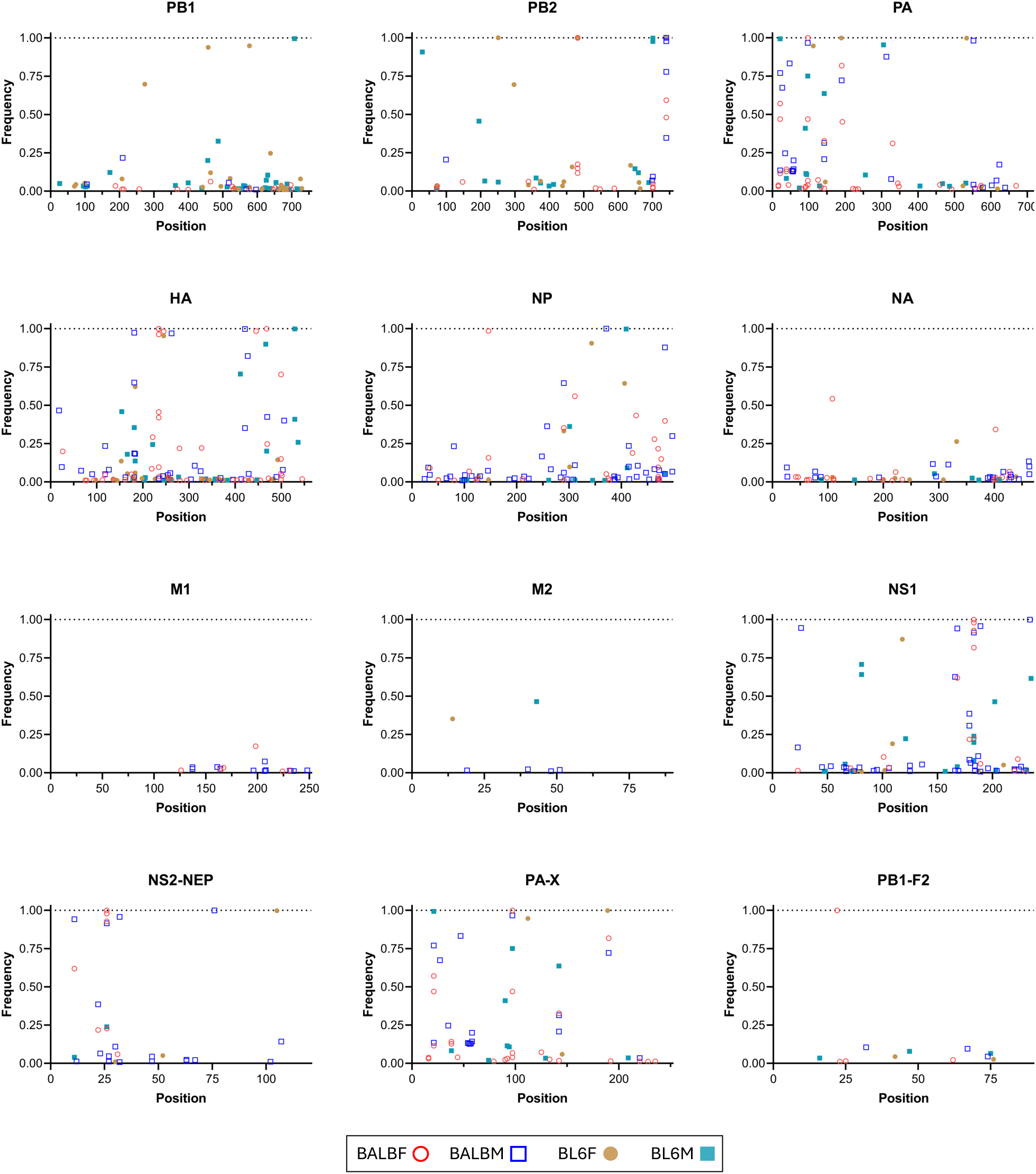
– Variant frequency maps for the 12 main influenza open reading frames. Variants are from 28 independent viral lineages, 7 lineages per host type.

**Table S1.** – Complete variant list for 28 host-adapted lineages. Available at https://github.com/RodrigoMamedeCosta/flu-evolution/ /Additional data/Variant list/TableS1 – Variant list.xlsx

**Table S2.** – Positively selected variants. Variant frequencies under each host are the mean of seven independent lineages. Bolded frequencies denote variants significant in the specific host at 6 Å.

| Protein | Variant | Freq<br>BALBF | Freq<br>BALBM | Freq<br>BL6F | Freq<br>BL6M | Reported effects | Refs |
| --- | --- | --- | --- | --- | --- | --- | --- |
| HA | N165D | 0.002 | **0.263** | 0.137 | 0.054 | Increased virulence | 20,21 |
| HA | G218W/E | **0.705** | 0.002 | - | - | Increased replication but not virulence. Increases fusion pH | 19,22 |
| NS1 | G183E/R | **0.567** | 0.131 | 0.016 | 0.074 | Interface-disrupting mutations can reduce virulence | 65,66 |
| PA | M21I | 0.157 | 0.129 | - | 0.142 | Increased polymerase activity | 22 |
| PA | T97I | 0.225 | 0.138 | - | 0.107 | Increased polymerase activity | 22 |
| PB2 | K482R | **0.349** | - | 0.143 | - | Increased polymerase activity, virulence and replication | 22 |
| PB2 | D701N | 0.014 | 0.014 | 0.129 | **0.425** | Increased polymerase activity, virulence and replication | 22 |
| PB2 | D740N | 0.160 | **0.332** | 0.143 | - | Increased polymerase activity, virulence and replication | 22 |

**Table S3.** – Deletion measurements and statistics. A) Comparisons between host types. B) Comparison between host strains. C) Comparisons between host sexes.

**A**
| Measure | BALBF | BALBM | BL6F | BL6M | KW statistic | p value |
| --- | --- | --- | --- | --- | --- | --- |
| Mean number of deletions | 3 | 4.2 | 5.43 | 8.29 | **40.53** | **<0.001** |
| Mean length of deletions | 7.25 | 14.71 | 19.16 | 22.22 | **11.23** | **0.011** |
| Mean frequency of deletions | 0.098 | 0.080 | 0.076 | 0.083 | 3.77 | 0.287 |

**B**
| Measure | BALB/c | C57BL/6 | W statistic | p value |
| --- | --- | --- | --- | --- |
| Mean number of deletions | 3.67 | 6.86 | **414.5** | **<0.001** |
| Mean length of deletions | 12 | 21.01 | **1156.5** | **0.021** |
| Mean frequency of deletions | 0.087 | 0.08 | 1682 | 0.599 |

| Measure | F | M | W statistic | p value |
| --- | --- | --- | --- | --- |
| Mean number of deletions | 4.55 | 6.58 | 2099.5 | 0.547 |
| Mean length of deletions | 16.3 | 20.23 | **1524** | **0.029** |
| Mean frequency of deletions | 0.081 | 0.083 | 1894 | 0.697 |

